# Applying PCR cycle autonormalization to improve PacBio full-length 16S rRNA sequencing

**DOI:** 10.64898/2026.01.13.699363

**Authors:** Charles J. Mason, Mikinley Weaver, Karma Kissinger, Melissa A. Johnson, Duan C. Copeland, Kirk E. Anderson, Scott M. Geib

## Abstract

The bacterial 16S rRNA gene is widely used to characterize host-associated and environmental microbiomes, most commonly through sequencing short hypervariable regions. Recent improvements in PacBio sequencing chemistry and concatenation approaches can now enable high-throughput, full-length 16S rRNA gene sequencing with high accuracy and depth. However, PCR-induced errors remain a major limitation, particularly for full-length amplicons where error accumulation may be elevated due to a five-fold increase in sequence lengths. These challenges are exacerbated when samples vary widely in microbial biomass, making it difficult to select a single optimal number of PCR cycles. Here, we evaluated the effects of PCR cycle autonormalization on PacBio Kinnex full-length 16S rRNA gene sequencing across seven agriculturally relevant specimen types. We compared conventional PCR protocols (20, 24, and 30 cycles) with an autonormalization approach in which individual reactions were terminated during exponential amplification based on real-time fluorescence thresholds. Autonormalized reactions consistently retained a higher proportion of sequences following denoising and chimera removal, exhibited fewer substitution-based errors, and produced more even read distributions across samples. Meanwhile, overamplified reactions (30 cycles) showed elevated error rates and higher sequence removal, particularly in samples with greatest microbial biodiversity. Importantly, PCR protocol had minimal effects on overall community composition relative to specimen types. These results demonstrate that PCR cycle autonormalization improves data yield for full-length 16S rRNA sequencing while maintaining robust inferences, while facilitating streamlined workflows for heterogeneous samples.

**Importance:** Amplicon-based sequencing of the 16S rRNA gene is foundational tool in microbiome research, yet PCR amplification remains a major source of technical error. This challenge is magnified for full-length 16S rRNA sequencing and for workflows that process specimen types with widely varying microbial biomass. Selecting a single PCR cycle number can result in under-amplification of samples with low 16S rRNA copies or over-amplification and error accumulation in high-titer samples. Here, we show that PCR cycle autonormalization, implemented using iconPCR, substantially improves PacBio full-length 16S rRNA sequencing by reducing PCR-induced substitution errors and chimeras while maintaining consistent biological inference. Autonormalization also enables blind pooling of amplicons without post-PCR quantification or equimolar normalization, reducing hands-on time and sample loss. These benefits make cycle autonormalization particularly valuable for high-throughput and production-scale sequencing applications handling diverse specimen types.

## Introduction

Amplicon-based sequencing of the bacterial 16S rRNA SSU (16S) gene has greatly advanced the study of microbial communities across host-associated and environmental systems (1, 2). Most microbiome studies rely on short-read sequencing of partial 16S regions generated using sequence-by-synthesis (SBS) chemistries, which provide high per base accuracy and scalability by taking advantage of the gene’s hypervariable regions for taxonomic classification (3–5). For some taxa, short fragments are insufficient to resolve closely related lineages, leading to ambiguous classification and reduced taxonomic precision (3, 6, 7). Sequencing near-full length 16S offers substantially improved taxonomic inferences and the potential to better resolve bacterial communities (8–16). Historically, adoption of long-read sequencing approaches has been constrained by lower read quality and throughput relative to short-read platforms. However, recent advances in PacBio circular consensus sequencing (CCS) have enabled the generation of highly accurate long reads (Q > 30, HiFi reads) (17, 18). In parallel, MAS-Seq concatenation strategies implemented through PacBio Kinnex protocols allow high-throughput full-length 16S sequencing at rates and costs comparable to traditional SBS based short-read amplicon sequencing (19). These developments enable hundreds to thousands of samples to be pooled into a single sequencing run, increasing the importance of consistent library preparation.

Upstream from sequencing, PCR is a major source of bias and error in amplicon-based analyses and can create artifacts that influence 16S sequence data (20–22). Polymerase choice, thermocycle number, and template concentration all influence substitution errors, chimera formation, and amplification bias (23–25). Excessive thermocycling can damage templates and propagate early errors into high-abundance artifacts that survive downstream quality filtering (23, 26–28). For full-length 16S sequencing, PCR-induced errors may be especially consequential because each read spans ∼5-7× more bases than short-read amplicons, increasing the probability that errors lead to sequence loss during QA/QC and processing into amplicon sequence variants (ASVs) or operational taxonomic units (OTUs).

Controlling PCR cycle number is particularly challenging when processing heterogeneous specimens. Microbial biomass and 16S gene copy number can vary by orders of magnitude between specimen types, making a fixed cycle number suboptimal for mixed sample sets (27, 29). Under-amplification reduces yield in low-biomass samples, while over-amplification increases error rates and chimera formation in high biomass samples. Having knowledge of sample composition can aid in mitigating cycling-related issues, but that can be challenging in scaled-production settings. Most workflows involve blind receipt and sample processing such that sample-specific protocols are unrealistic.

In this study, we evaluated the use of PCR cycle autonormalization using iconPCR (where individual reactions are automatically terminated during exponential amplification based on real-time fluorescence thresholds) (28) to improve full-length 16S microbiome analysis. Using agriculturally-relevant arthropod, plant, and soil specimens with varying microbial diversity and titers, we compared autonormalized PCR to defined-cycle protocols across multiple metrics of sequence quality, retention, and community inference. We anticipated that excessive PCR cycling would increase error-driven sequence loss, disproportionately affect diverse specimen samples, and that cycle autonormalization would mitigate these effects while preserving biological signal.

## Methods

### Samples

Seven environmental specimen types were selected to represent a wide range of microbial richness, diversity, composition, and dominant taxonomic groups (Supplemental Table 1). Samples included five arthropod species from distinct orders (pickleworm – *Diaphania nitidalis* - Lepidoptera; *Lema* beetle - *Lema sp.* - Coleoptera; termite - *Cryptotermes brevis* – Blattodea; honey bee - *Apis mellifera* - Hymenoptera; and medfly - *Ceratitis capitata* - Diptera), coffee (*Coffea arabica var. typica*) leaf phyllosphere swabs, and soil collected at the base of coffee trees. Environmental sample sizes ranged from 6-8 replicates per group. DNA extraction methods varied among samples because materials were obtained from independent research projects (Supplemental Table 1). We used commercial positive controls alongside experimental samples, including two DNA standards (2 ng per reaction, Zymobiomics Microbial DNA Standard Lot ZRC193007 Zymo Research Corp, Irvine, CA, USA; henceforth ‘DNA control 1 & 2’) and one community standard processed through DNA extraction (Zymobiomics Microbial Community Standard Lot ZCR213683; henceforth ‘extraction control’).

### PCR amplification procedures

PCRs were performed using the iconPCR instrument (n6 Tec, Pleasanton, CA, USA) (28). The iconPCR instrument possesses individual well control such that amplification cycles may be ceased on a well-by-well basis following a designated change in fluorescence via intercalating dyes. Reactions (15 µL) included 0.25 µM of dual-indexed 27F and 1492R primers contained in-line 10-mer barcodes and indices for concatenation PCR asdescribed in the PacBio Kinnex 16S Kit (Pacific Biosciences, Menlo Park, CA, USA). Q5 polymerase was chosen over other options because in preliminary optimization of amplification protocols, there was minimal incidence of primer dimer formation and false positive signal on the iconPCR instrument compared to other hi-fidelity polymerases (Supplemental Figure 1).

Number of PCR cycles were varied to assess the impact of autonormalization vs. predetermined cycle number (Figure 1). PCR cycling conditions were 98 ℃ for 30s, variable numbers of cycles consisting of 98 ℃ for 10s, 57 ℃ for 15s, 72 ℃ for 2 min, followed by a final extension at 72 ℃ for 10 min. For autonormalization procedures, cycling was terminated by changes in the slope of the samples’ fluorescence, with maximum threshold set at 30 cycles. This technique employed the iconPCR software ‘Slope mode’ which terminates cycles in the exponential phase of amplification prior to plateauing (Supplemental Figure 2; cycle terminations reported in Supplemental Table 1). Fixed-cycle protocols of 20, 24 and 30 cycles were performed for comparison. All protocols were performed on the iconPCR instrument.

**Figure 1:**
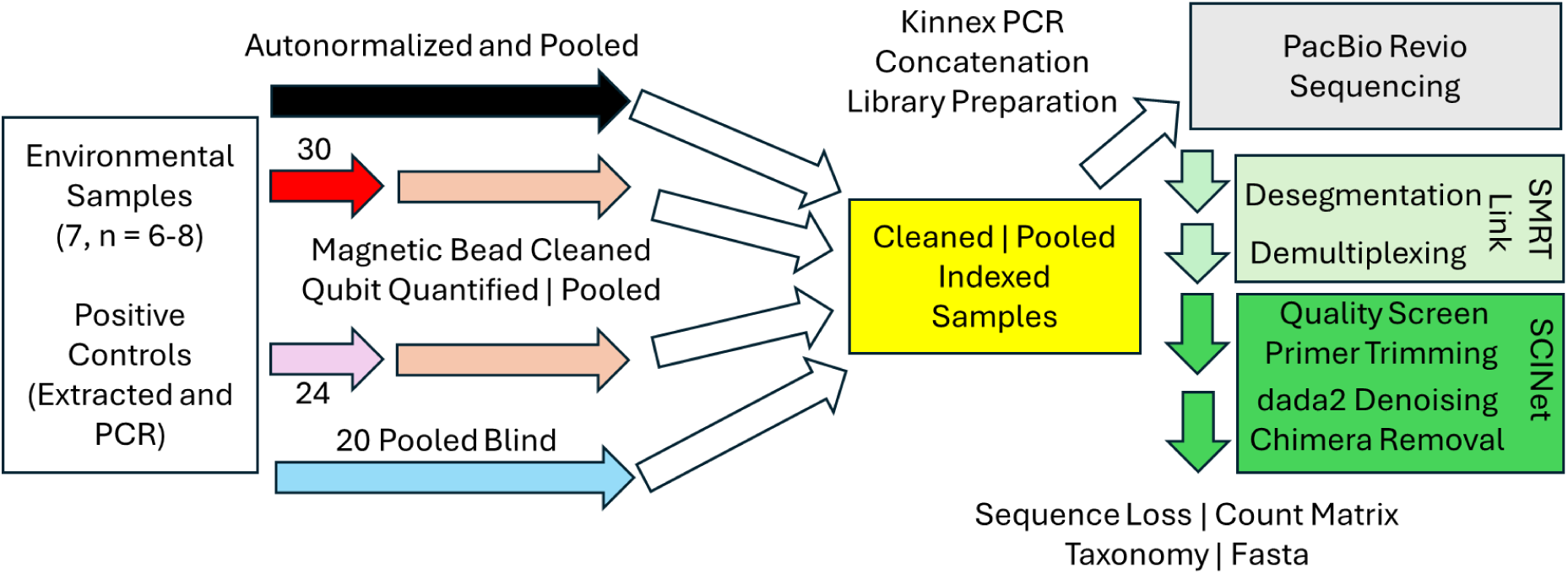
Schematic of experimental workflow. Cycling protocols varied between autonormalization, and *a priori* designated protocols of 30, 24, and 20 cycles. Samples were barcoded using dual-indices and pooler prior to concatenation and sequencing of reads. Specimens tested were included to provide a range in putative bacterial biodiversity and were agriculturally relevant. We performed no upstream QA/QC of sample types to simulate blind sample receipt and processing.

### Pooling, library preparation, and sequencing

Following amplification, 5 µL of each reaction was pooled for the autonormalized protocol and 20-cycle protocol and were purified using AMPure bead and quantified with a fluorometer (DeNovix Inc., Wilmington, DE, USA). For the 24- and 30-cycle protocols, samples were bead purified, quantified, and pooled in equimolar concentrations to reflect common normalization practices. Following pool quantification, amplicon libraries were combined into a single pool for Kinnex PCR and concatenation procedures in equivalent DNA concentrations.

Kinnex followed manufacturer kit protocols, except incubation times were doubled. The ∼19 kb concatenated amplicon library was sequenced on a single Revio 24M SMRT cell on a Pacific Biosciences Revio system. HiFi reads resulting from the run were desegmented and demultiplexed using PacBio SMRTLink v.13 before further processing.

### Sequence processing

HiFi reads were filtered to retain reads with mean read quality score ≥Q30 and per-base quality score ≥Q20 using fastplong v.0.4.1 (30). For per-base quality score, we designated a 2% cutoff rate where if low quality base scores exceeded that value in the read the sequence was culled. We then discarded sequences by length (min = 1300; max = 1550) before using cutadapt v.5.1 (31) to trim primers and reorient reads into the same direction. Reads lacking both forward and reverse primers were removed from the dataset. Amplicon sequence variants (ASVs) were generated with DADA2 in R v.4.4 (32, 33). Reads were filtered with DADA2 with expected error = 2 (34), dereplicated, and the DADA2 error learning algorithm adjusted for binned quality scores was used to infer error rates (PacBio Revio quality scores: 3, 10, 17, 22, 27, 35, 40). After estimating error rates, DADA2 denoising algorithms were used to infer sequences and chimeric sequences were removed using dada2::isBimeraDenovo using pooled samples. ASVs were taxonomically classified using the Naïve Bayesian Classifier implemented in DADA2 against the SILVA database v.138.2 (35–37). Control error rates before and after processing were conducted with .fastq files that were base quality and size filtered using mothur v.1.48.3 with the seq.error command (38). After DADA2 processing, ASV count and fasta file were converted into mothur formats for end-pipeline processing to determine control error rates under different conditions. We used manufacturer provided supplied reference sequences that matched the control lots.

### Data analysis

Statistical analyses were performed using R v4.4 (39). Alpha-biodiversity metrics were computed using the R package vegan (40) and averaged across different rarefaction events to accommodate differing degrees of sampling intensity (10,000 reads subsampled, 100 iterations). Relationships between sample Shannon H‵ and sequence retention were performed using Spearman correlation coefficients using the R package rstatix (41). PCR cycles and alpha diversity metrics were compared using pairwise Wilcox rank-sum tests with FDR correction. Beta-diversity indices were computed following rarefaction (100 iterations). For Bray-Curtis, rarefaction iterations were performed using avgdist in vegan with a subsample cutoff of 10,000 reads. For Jaccard, samples with counts below 10,000 were removed before repeated sampling and computation of the dissimilarity matrix. Non-metric multidimensional scaling was performed on the resulting matrices to determine separation of samples. PERMANOVA analyses were conducted using vegan::adonis2 with sample type and PCR protocols as fixed effects, and a strata subgroup effect was included on each individual sample type to account for sample-to-sample variability. Pairwise PERMANOVA was performed using the package pairwiseAdonis (42).

## Results

### Sequencing output following initial processing

The sequenced library produced 3.6M HiFi reads and 63.6 Gb of raw sequence data, with 94.3% of reads having an average base quality ≥ Q30. Mean and median length of concatenated reads were 17.9kb and 18.9kb, respectively. Desegmentation yielded 37,699,404 16S reads with an average length of 1,554 bp. Autonormalization of libraries generated the largest number of useable reads after quality control compared with fixed-cycle protocols (11.3M for autonormalization, 8.9M for 30 cycles of PCR, 7.3M for 24 cycles of PCR, and 8.1M for 20 cycles of PCR) (Figure 2).

**Figure 2:**
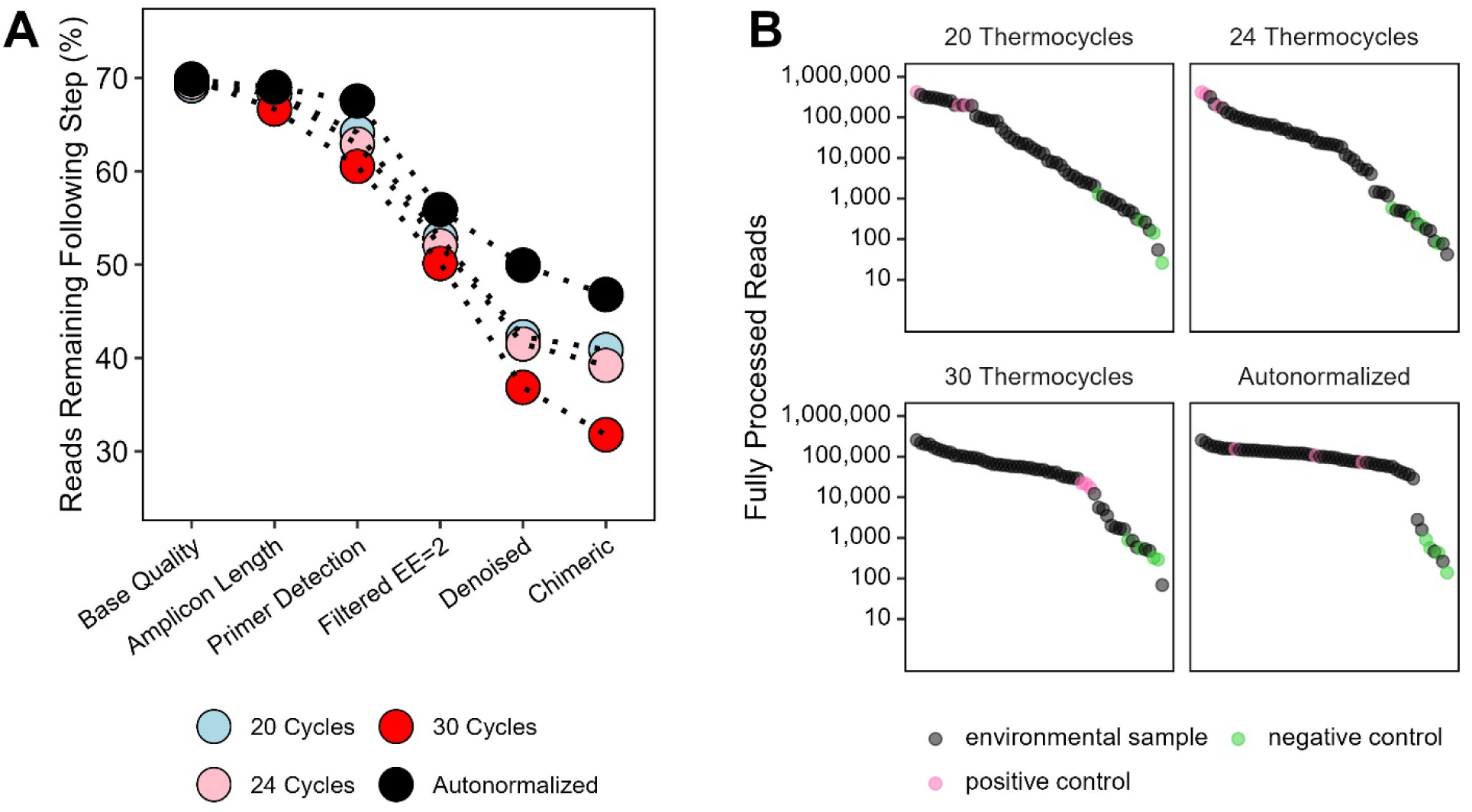
PCR cycles have differential impacts on sequenced libraries. (A) Thermocycle protocol exhibited different levels of sequence loss at the different filter steps. All protocols exhibited approximately the same sequence loss (∼30%) due to low quality bases (>2% at <Q20). Higher (30) PCR cycles had more sequences culled with dada2 denoising and chimera checking protocols compared to the other PCR protocols. (B) Distribution of number of starting (upper point) end-process (lower point) reads between different PCR protocols. Data plotted on a log_10_(*y*) scale and samples are distributed by rank-abundance of the final reads along the x-axis.

### Effects of PCR protocol on sequence retention

PCR cycle number strongly influenced sequence retention during denoising and chimera removal. While all protocols lost ∼30% of reads during base-quality filtering, samples amplified with 30 cycles experienced substantially greater losses during denoising compared to autonormalized reactions. The 20- and 24-cycle procedures had sequence retention percentages between that of the autonormalization and 30-cycle protocols (Figure 2A). After fully processing reads, the number of sequences that were retained varied between protocols (Figure 2B). The 20-and 24-cycle protocols had the greatest spread in the number of reads per environmental sample, while the 30-cycle protocol was more even in distribution. Autonormalization consistently produced the most even distribution of reads across samples following full processing.

### Specimen-specific effects of PCR cycle

Sequence loss varied by specimen type and PCR protocol (Figure 3). Samples with high biodiversity, particularly soils, were disproportionately affected by overamplification, where <10% of reads were retained at 30 cycles. Given that soil samples had much higher biodiversity (Figure 4, Supplemental Table 3), we ran additional correlations without the samples but found that the correlations were largely unchanged except for the 20-cycle protocol (p = 0.078). Some arthropod samples (honeybee, termite, and pickleworm) had higher percentages of sequences filtered at amplicon length and primer detection step compared to others. Across protocols, sequence retention was negatively correlated with Shannon H‵ (20: σ = −0.33, p = 0.021; 24: σ = −0.61, p < 0.001; 30: σ = −0.78, p < 0.001; autonormalized: σ = −0.66, p < 0.001) (Figure 4).

**Figure 3:**
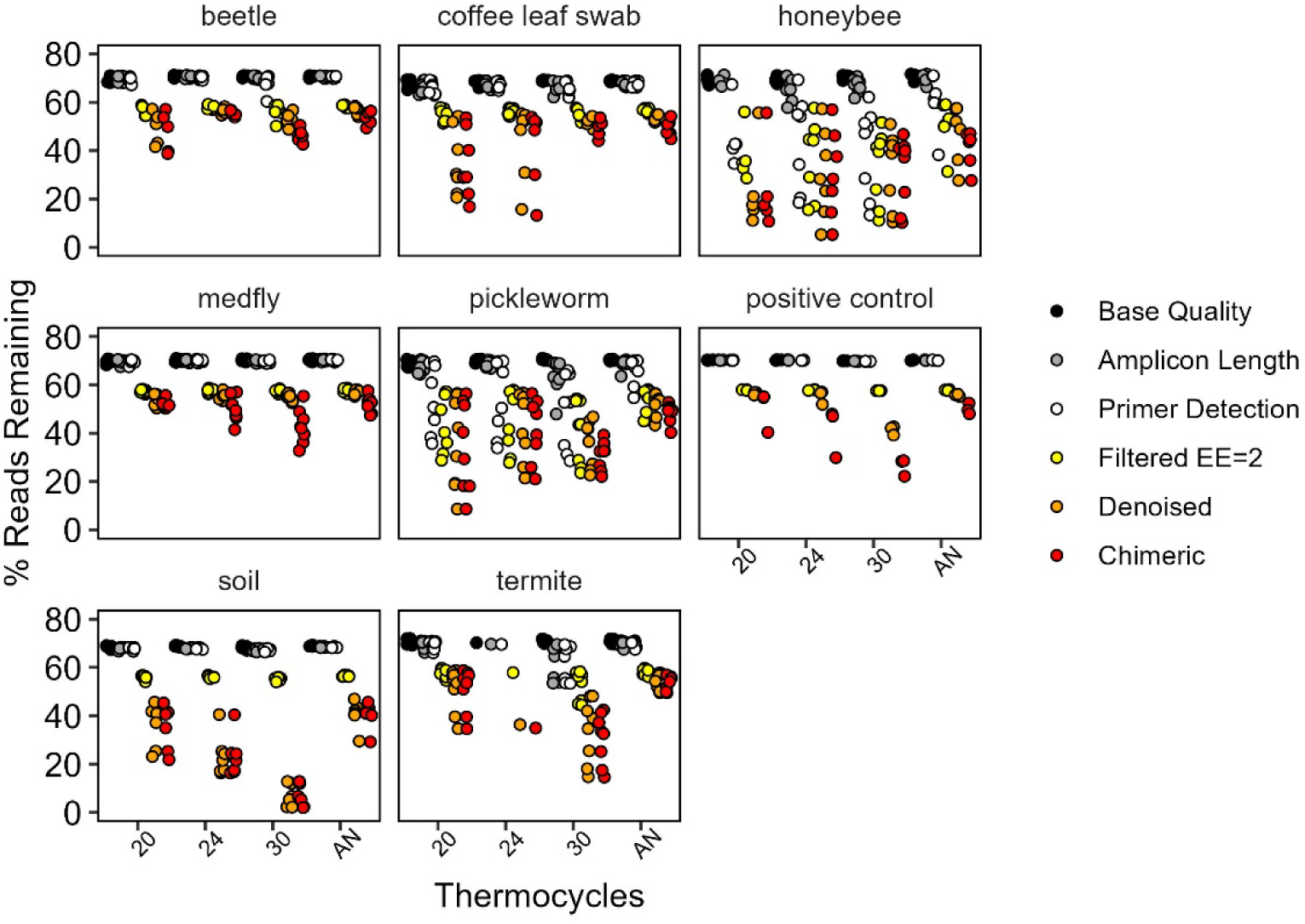
PCR protocol effects on retained reads (%) after filtering steps for different specimen types. Only samples with ≥3000 sequenced reads are displayed. AN = autonormalization of PCR thermocycles.

**Figure 4:**
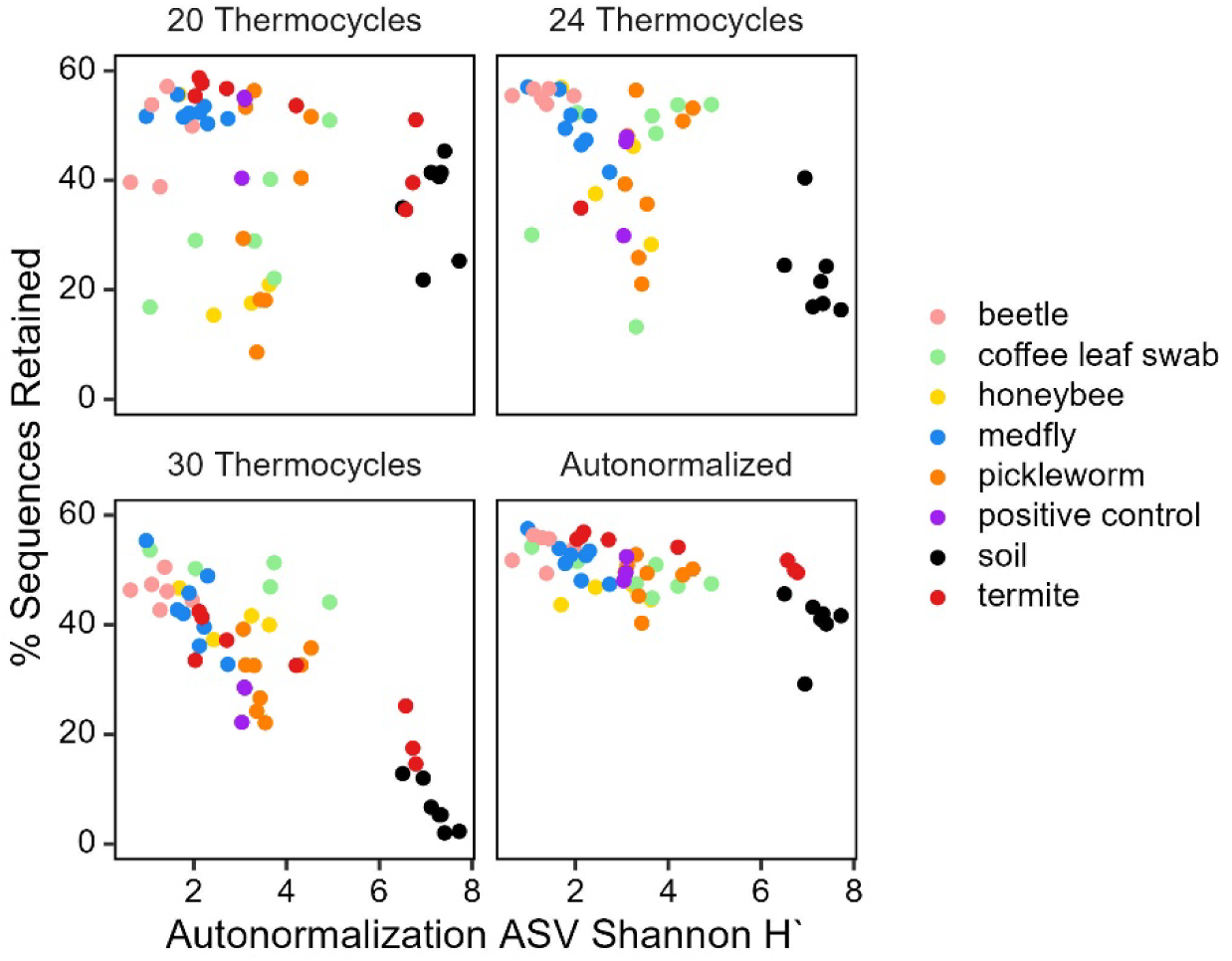
Relationships between sample ASV alpha-diversity and sequence loss. Generally, with higher Shannon diversity had fewer sequences retained, with thermocycle procedures exacerbating the retention of samples. Only samples with sequences ≥3000 at start of process are displayed. Shannon H‵ was computed using 10,000 sequences rarefied and averaged for 100 iterations using autonormalization PCR methods.

### Error profiles in positive controls

PCR protocol had minimal effects on overall taxonomic composition of positive controls (Figure 5A). However, 30-cycle reactions exhibited elevated Shannon diversity (Figure 5B) and ASV richness (adjusted for number of reads) (Figure 5C), consistent with spurious sequence inflation (p < 0.010; Supplemental Table 3). Heatmap of the two DNA samples indicated strong clustering between the technical replicates between PCR protocols (Figure 5D).

**Figure 5:**
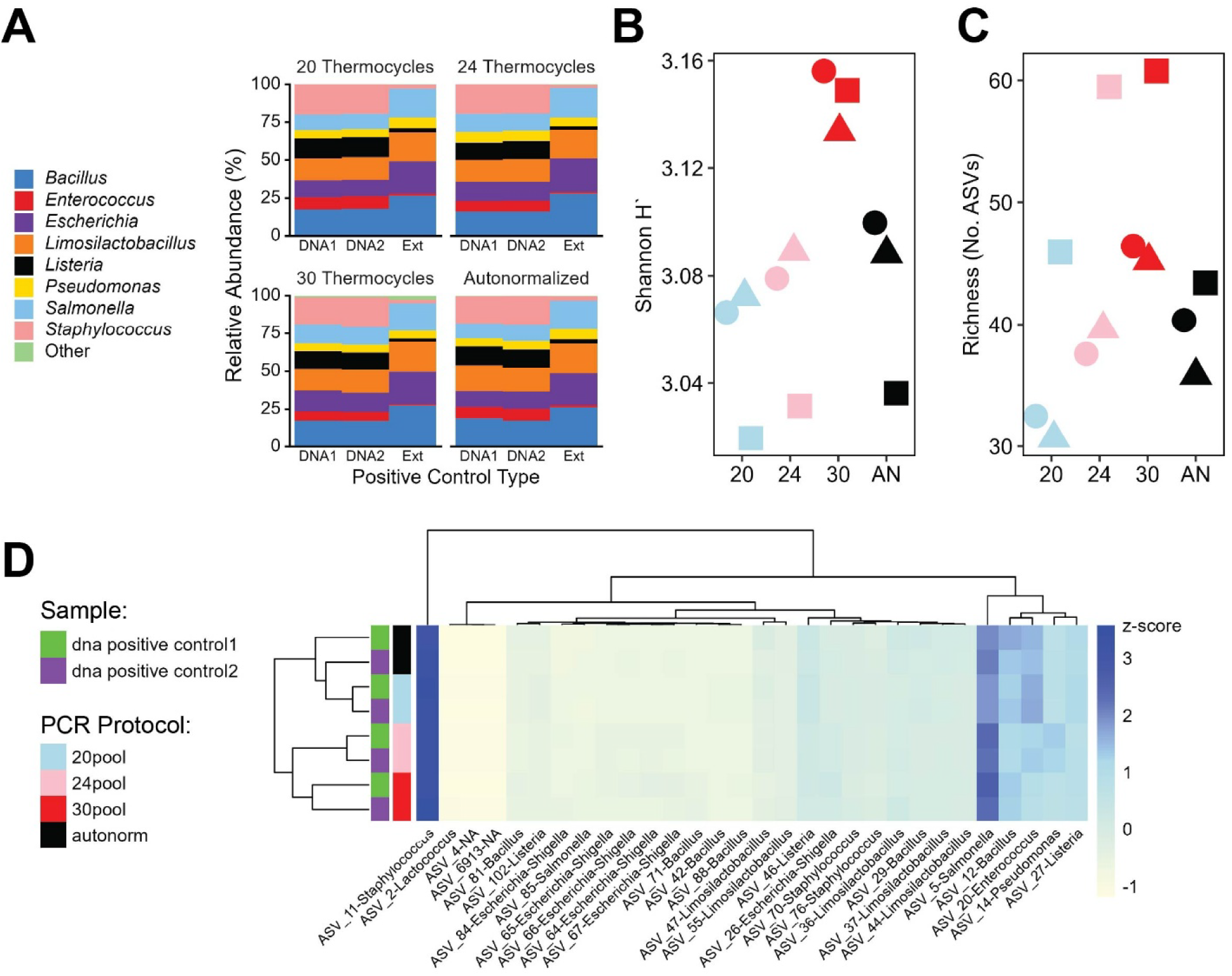
Impact of thermocycles on positive controls. (A) At the genus sequence level, there were few differences in the ASV composition, except some elevated ‘other’ sequences in the extracted controls. Theoretical relative abundance of controls: *Bacillus –* 17.4%; *Staphylococcus* 15.5%; *Salmonella* 10.3%; *Escherichia* 10.1%; *Listeria* 14.1%; *Limosilactobacillus* 18.5%; *Enterococcus* 9.9%; and *Pseudomonas* 4.1% (B) Shannon diversity indices and (C) the number of ASVs adjusted for the number of sample sequence were elevated in control samples receiving 30 PCR cycles compared to other procedures. Shapes denote the different controls amplified (circle = DNA Positive 1, triangle = DNA Positive 2, square = Extracted).

Substitution-based errors dominated error profiles and increased with PCR cycle number (Figure 6A), with non-random positional distributions along the 16S gene (Figure 6B). Error rates for DNA controls after processing were 3.3-3.6 × 10^-4^ for autonormalization and 20 and 24-cycle protocols and 3.9-4.9 × 10^-4^ for 30-cycle methods (Supplemental Table 4). Manual inspection of controls indicated some likely laboratory cross-contaminants at low sequence relative abundances (< 0.001%), which adversely affect the error rates (Supplemental Table 5). Several taxonomic matches had consistent mismatches between the references across the control technical replicates (Supplemental Table 5), which would indicate that these were true sequence variants rather than one-off PCR errors.

**Figure 6:**
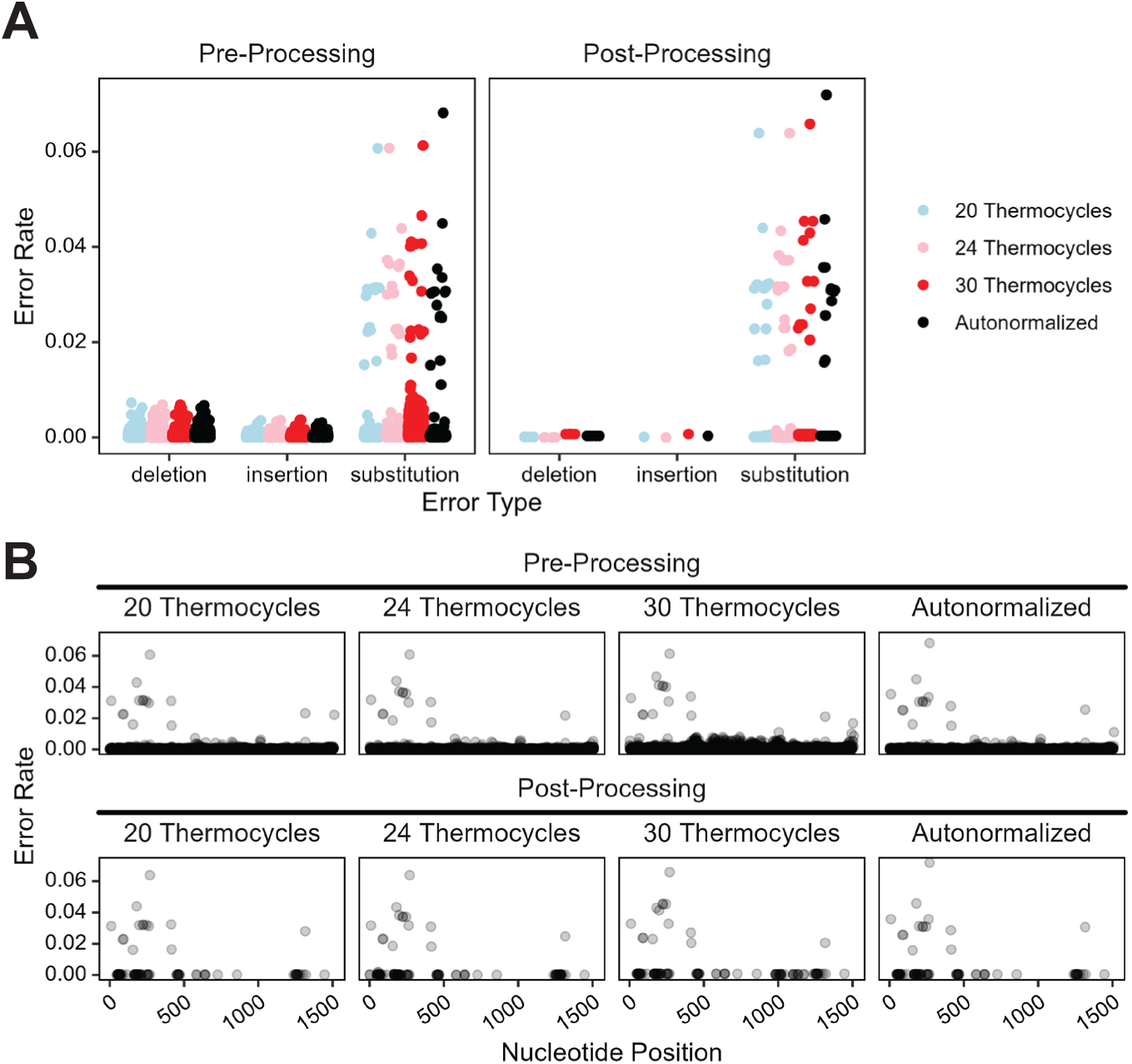
Impact of thermocycles on error rate of positive control (DNA Positive 1). Error rates were computed by the sum of mismatches to reference / sum of bases in query before and after QA/QC processing through DADA2. (A) There was a greater incidence of substitution-based errors across the dataset, with greater numbers being present in the samples having greater PCR cycles. (B) Positional incidence of all errors indicate some consistency in where the mismatches to references were located across the sequences. Substitution error rates for other positive controls are provided in Supplemental Figure 3.

### Community composition and diversity metrics

Ordination clustered samples primarily by specimen type rather than PCR protocol for both Bray-Curtis and Jaccard computed distances (Figure 7). PERMANOVA indicated that PCR protocol explained a small but statistically significant fraction of variation (R^2^ < 0.06), whereas specimen type accounted for the majority of variance (R^2^ > 0.27; Table 1). There was a high degree of residual sum of squares for both PERMANOVA models, suggesting a much of the unexplained variance that are likely attributed to individual samples. Alpha-diversity metrics were largely consistent across protocols (Figure 6C & 6D; Supplemental Table 5), with limited specimen-specific differences. Termite samples had greater Shannon metrics (p = 0.03) and ASV richness (p = 0.016) in the autonormalized reactions compared with 30 cycles.

**Figure 7:**
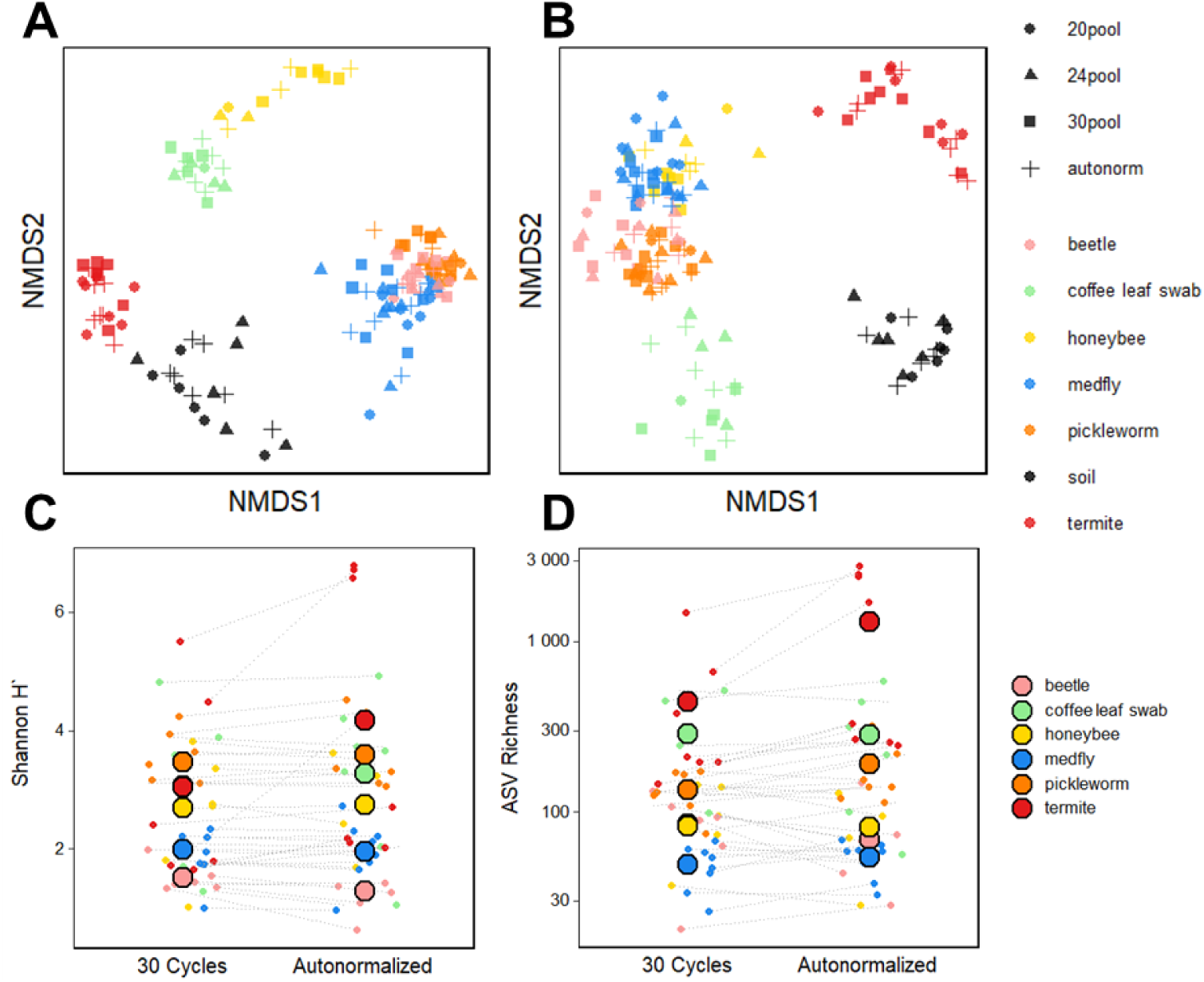
Ordination and biodiversity metrics of environmental samples. (A) NMDS using Bray-Curtis dissimilarities indicate strong clustering by sample type, with little overt impact of PCR protocol. (B) NMDS of specimens using Jaccard (presence/absence) indicates a first grouping of samples. (C) Shannon diversity and (D) ASV richness of environmental samples at 30 cycles vs. autornormalzied protocols. Large circles represent the mean within each group of specimens. For both alpha- and beta-biodiversity metrics, soil samples did not obtain enough processed read counts (less than 10,000) to adquate comparisons at 30 PCR cycles with other metrics.

**Table 1:** PERMANOVA results of environmental specimens in response to different PCR protocols. Models were built with individual sample included as a ‘strata’ effect.

A. Bray-Curtis Dissimilarity:
|  | df | MSS | R<sup>2</sup> | F-value | *p-value* |
| --- | --- | --- | --- | --- | --- |
| Sample Type | 6 | 27.3 | 0.39 | 14.0 | 0.001 |
| PCR Protocol | 3 | 0.26 | 0.004 | 0.26 | 0.001 |
| Sample Type * PCR Protocol | 16 | 1.79 | 0.02 | 0.34 | 0.001 |
| Residual | 123 | 49.8 | 0.58 |  |  |
| Total | 148 | 70.1 | 1 |  |  |

B. Jaccard Dissimilarity:
|  | df | MSS | R<sup>2</sup> | F-value | *p-value* |
| --- | --- | --- | --- | --- | --- |
| Sample Type | 6 | 16.9 | 0.24 | 7.27 | 0.001 |
| PCR Protocol | 3 | 0.85 | 0.01 | 0.73 | 0.001 |
| Sample Type * PCR Protocol | 16 | 4.44 | 0.06 | 0.72 | 0.002 |
| Residual | 123 | 47.7 | 0.68 |  |  |
| Total | 150 | 69.9 | 1 |  |  |

## Discussion

Long-read sequencing techniques and their application to microbiome or 16S rRNA amplicon sequencing have been hampered in part by poor read quality and sequencing yields. Advances in PacBio sequencing chemistry coupled with MAS-Seq concatenation strategies have made high-throughput, full-length 16S sequencing increasingly accessible. Our study produced read output and quality commonly employed SBS techniques for hypervariable regions. However, PCR remains a critical point in processing, particularly for workflows handling heterogeneous specimens to be combined into a single sequencing pool. In this study, we demonstrate that PCR cycle autonormalization substantially improves data retention and reduces PCR-induced errors without altering overall biological inference. Higher number of PCR cycles consistently increased sequence loss during denoising and chimera removal, especially in the more biodiverse and higher-titer samples. Because denoising algorithms infer true sequences based on abundance and error models (8, 32), substitution errors introduced early in PCR can propagate into abundant but erroneous variants that are subsequently removed. This effect is magnified for full-length amplicons, where even low per-base error rates increase the probability of read rejection through processing steps.

Substitution-based errors were the dominant error type and increased with PCR cycle number, consistent with other studies sequencing mock communities (43, 44). The non-random distribution of errors suggests systematic amplification-associated mechanisms rather than stochastic sequencing noise (43). The reference fasta files provided with the commercial mock communities indicated 48 16S entries from the eight known bacteria in the references, and as others have noted the reference is potentially incomplete and/or incorrect (8, 45). Like these other studies, amplicons with mismatches with the reference were identified by other classifiers (8, 45). In our manual inspection, there are low abundance reads arising from other samples occurring in very low abundances, but we opted to retain these for our determination of the error rate since they represent likely scenarios to occur during the processing of these types of samples. Contamination could potentially arise from laboratory contamination during PCR setup. One additional aspect that may contribute to sample contamination of may be related to the Kinnex library preparation and potential chimeric read formation or barcode switching or at demultiplexing steps. For example, we observed sporadic, inconsistent incidences of these dominant taxa also present in negative controls. This an important limitation related to Kinnex approaches and its application to low-abundant taxa, but could be ameliorated with unique molecular identifier-based methods (45, 46). While absolute error estimates may be influenced by limitations of commercial reference sequences and potential laboratory contaminants, relative comparisons across PCR protocols indicate that autonormalization did reduce substitutions introduced during PCR.

PCR protocol had a minimal impact on community-level inferences. Although statistically significant effects were detected, effect sizes were small relative to specimen-specific variation, indicating that autonormalization improves technical performance without distorting ecological conclusions. These findings align with previous short-read studies showing limited compositional effects of PCR cycle number under controlled conditions (25, 27, 47, 48). However, others have indicated that overamplification can impact rare amplicons in diverse samples (28, 49), so the specific impacts of thermocycling protocols may be case-by-case or sample-by-sample. With exception of the soil specimens, our samples had relatively low microbial biodiversity compared with that of other environmental sample types. So, although the effect of PCR protocol was statistically significant in our sample types, the lack of a strong difference may be due to the type of specimens that we used and the fact that the majority of the high diversity samples were culled in the sequence processing.

The fact that more biodiverse samples were more susceptible to sequence loss likely results from combination of factors. First, the most biodiverse sample we used also possessed the greatest titers (soil), therefore these samples received upwards of eleven extra cycles to introduce additional errors compared to the autonormalization protocol. Second, a greater amplicon biodiversity may produce a greater number of erroneous samples. Regardless of the specific mechanism, these observations are important considerations for tuning PCR protocols for putatively speciose specimens.

Currently, there are few specific protocols or best practices for handling full-length 16S data and the technical aspects of data quality control were outside of the specific goals of this study. The processing steps we utilized can be modified and shifted depending on the specific objectives. However, in the initial data curation we found that with relaxing upstream processing steps for soil samples, the majority were still culled at denoising steps indicating that there were unrecoverable issues with the sequenced data. Similarly, in some of the arthropod specimens we observed that a greater number were removed due to amplicon lengths, which appeared to be related to non-specific priming of the host 18S. For downstream data analysis, production of full-length 16S amplicons has a high likelihood that same bacteria is split into different ASVs (8, 50, 51), so clustering of ASVs into operational taxonomic units and a 3% or 1% cutoff may be preferred for certain applications (52, 53). Additional steps involving statistical programs to handle errant laboratory contaminants may also aid in the inferences of these datasets (54).

## Conclusions

Full-length 16S gene sequencing offers advantages for taxonomic resolution and phylogenetic inference, but its broader adoption depends on reliable and scalable upstream laboratory workflows. When amplifying of specimens with variable microbial biomass, determining the appropriate number of thermocycles for PCR protocols can be challenging. Samples with lower microbial titers may be under amplified, while samples with excessive cycles can incur high error rates and reduce the data quality. The individual well control like that of the iconPCR instrument helps to mitigate both issues by tightly controlling the number of thermocycles for specific samples. Our results demonstrate that PCR cycle autonormalization improves performance of PacBio full-length 16S sequencing by reducing PCR-induced substitution errors, improving read retention, and producing more even sample representation across heterogeneous sample types, eliminating the need for post-PCR normalization. Importantly, this approach enhances both data quality and laboratory efficiency, particularly for production-scale microbiome sequencing.

## Contributions

CJM designed experiments, collected samples, performed PCR, analyzed data, and wrote the first draft of manuscript; MW, KW, MAJ, DCC, and KEA provided samples for PCR and analysis; SMG designed experiments, provided feedback on data analysis, and provided editorial feedback. All coauthors approved of the manuscript before submission.

## Acknowledgements

We thank Angela Kauwe and Nicholas Newell for technical assistance for final library preparation and sequencing. The findings and conclusions in this publication are those of the authors and should not be construed to represent any official USDA or U.S. Government determination or policy. Mention of trade names or commercial products in this publication is solely for the purpose of providing specific information and does not imply recommendation or endorsement by the U.S. Department of Agriculture. USDA is an equal opportunity provider and employer. This research used resources provided by the SCINet project and/or the AI Center of Excellence of the USDA Agricultural Research Service, ARS project numbers 0201-88888-003-000D and 0201-88888-002-000D. This research was supported by the U.S. Department of Agriculture, Agricultural Research Service appropriated project “Advancing Molecular Pest Management, Diagnostics, and Eradication of Fruit Flies and Invasive Species” (2040-22430-028-000-D).

---

**Supplemental Figure 1:**
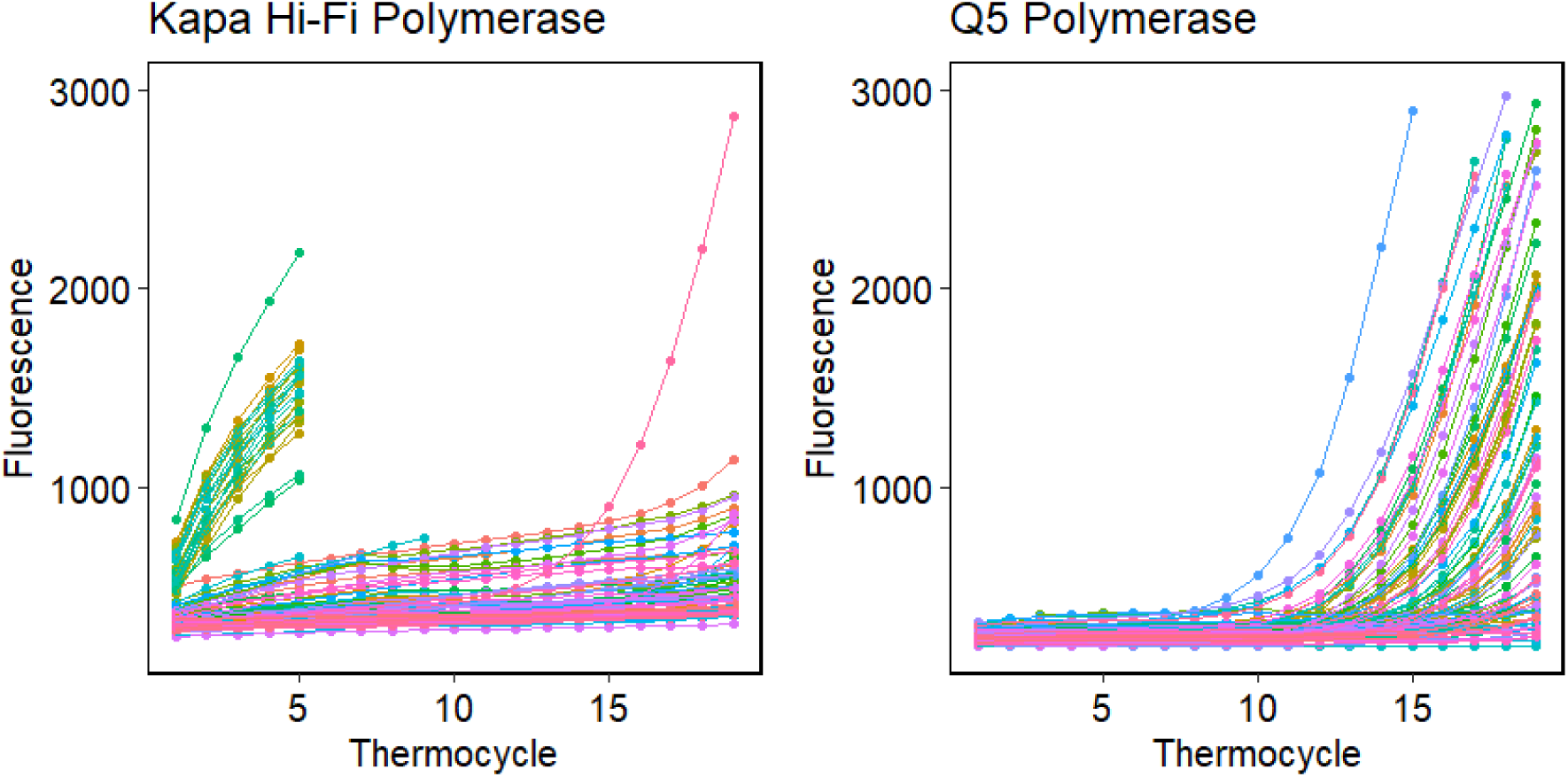
Fluorescence signal comparison between two polymerases with same 16S barcoded primer set. Note early terminations in left panel indicative of the formation of primer dimer.

**Supplemental Figure 2:**
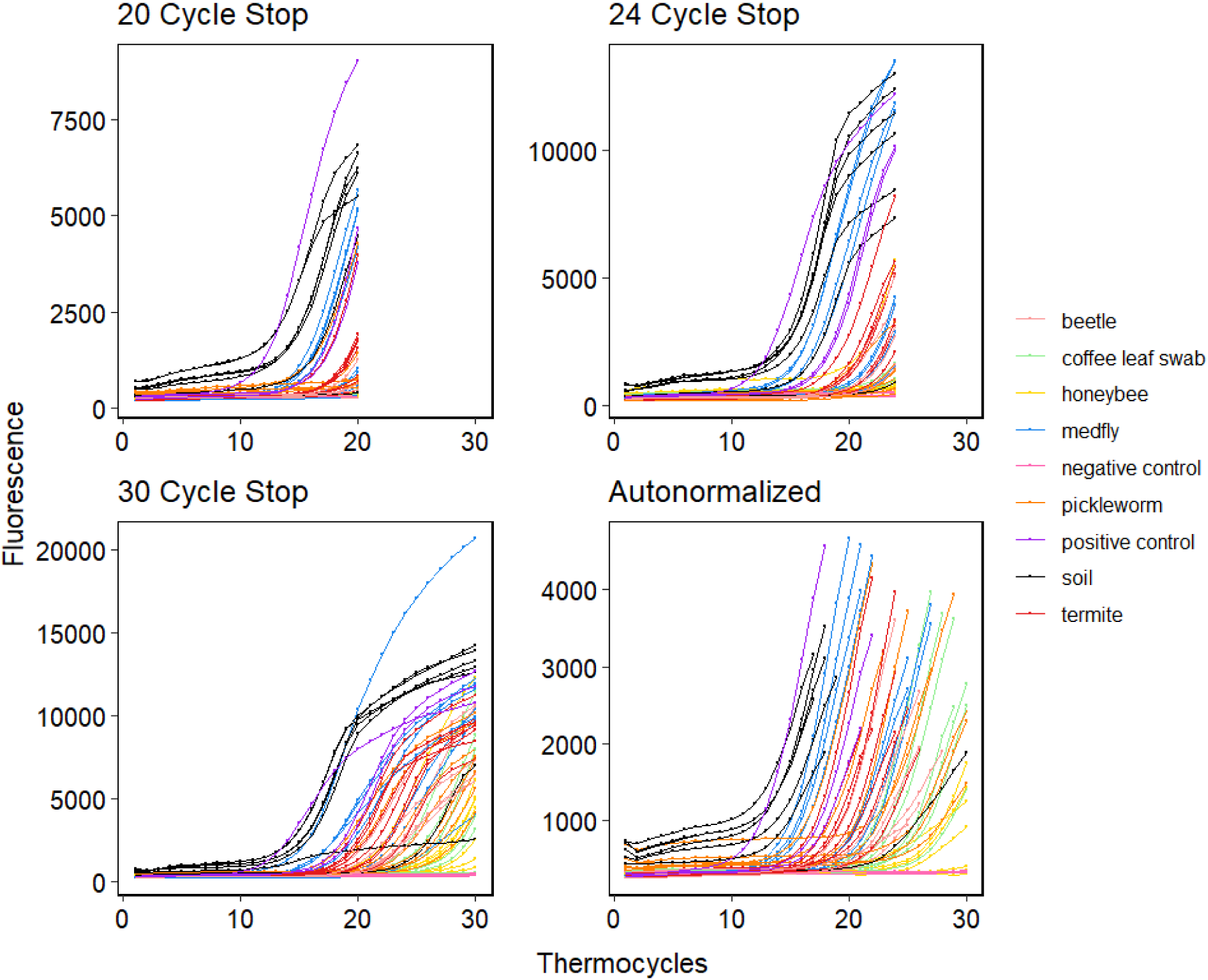
Fluorescence signal of environmental and control systems under different thermocycler protocols.

**Supplemental Figure 3:**
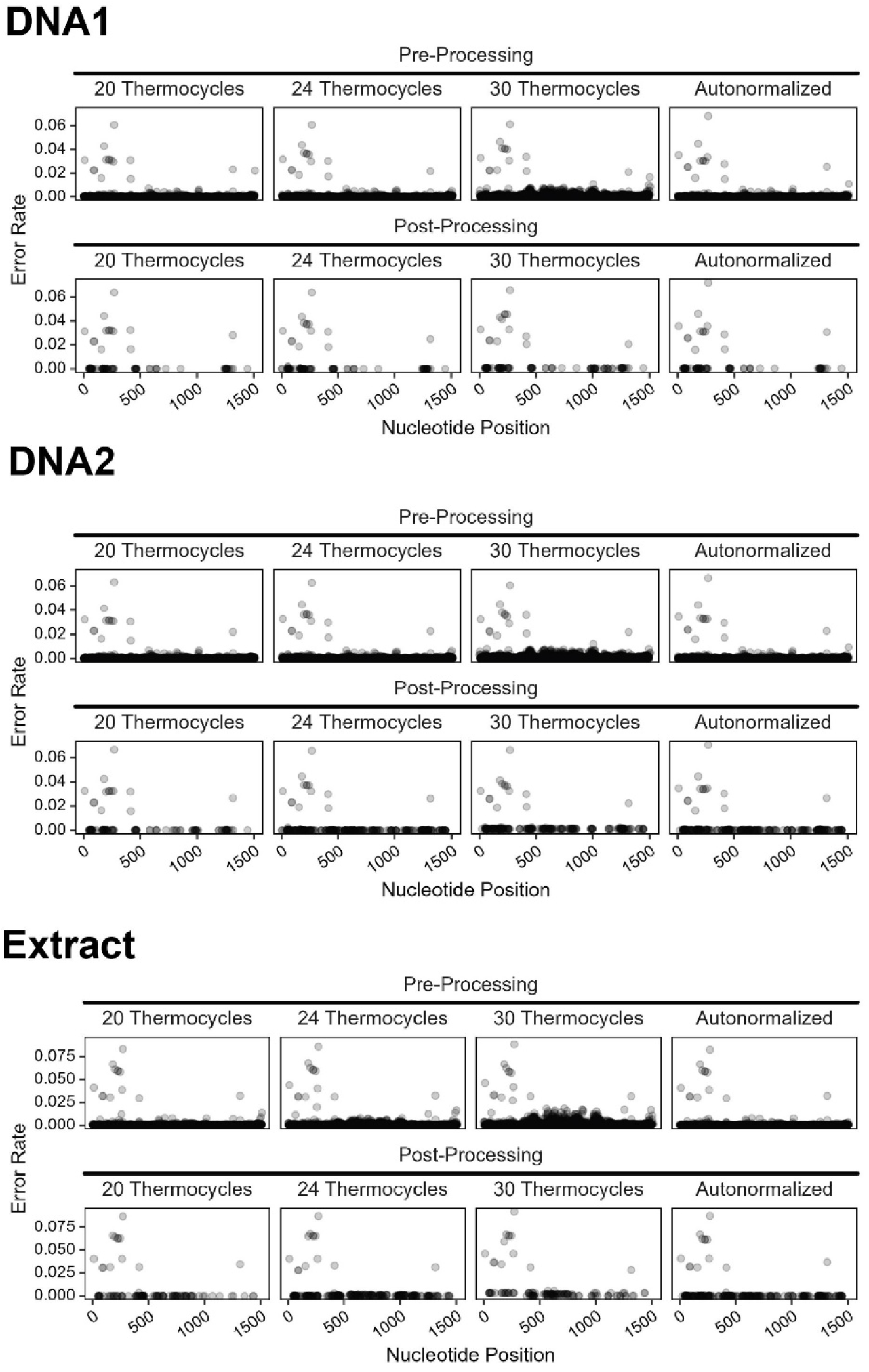
Substitution-based error rates of positive control samples.

